# Causal associations between risk factors and common diseases inferred from GWAS summary data

**DOI:** 10.1101/168674

**Authors:** Zhihong Zhu, Zhili Zheng, Futao Zhang, Yang Wu, Maciej Trzaskowski, Robert Maier, Matthew R. Robinson, John J. McGrath, Peter M. Visscher, Naomi R. Wray, Jian Yang

## Abstract

Health risk factors such as body mass index (BMI), serum cholesterol and blood pressure are associated with many common diseases. It often remains unclear whether the risk factors are cause or consequence of disease, or whether the associations are the result of confounding. Genetic methods are useful to infer causality because genetic variants are present from birth and therefore unlikely to be confounded with environmental factors. We develop and apply a method (GSMR) that performs a multi-SNP Mendelian Randomization analysis using summary-level data from large genome-wide association studies (sample sizes of up to 405,072) to test the causal associations of BMI, waist-to-hip ratio, serum cholesterols, blood pressures, height and years of schooling (EduYears) with a range of common diseases. We identify a number of causal associations including a protective effect of LDL-cholesterol against type-2 diabetes (T2D) that might explain the side effects of statins on T2D, a protective effect of EduYears against Alzheimer’s disease, and bidirectional associations with opposite effects (e.g. higher BMI increases the risk of T2D but the effect T2D of BMI is negative). HDL-cholesterol has a significant risk effect on age-related macular degeneration, and the effect size remains significant accounting for the other risk factors. Our study develops powerful tools to integrate summary data from large studies to infer causality, and provides important candidates to be prioritized for further studies in medical research and for drug discovery.

## Introduction

Health risk factors such as body mass index (BMI), serum cholesterol and blood pressure are associated with many human common diseases ^1,2^, e.g. being overweight is associated with increased risk to cardiovascular diseases (CVD) ^3^ and type-2 diabetes (T2D) ^4^. These associations are usually derived from observational studies that cannot distinguish whether the risk factors are ‘upstream’ causal factors, ‘downstream’ consequences of the diseases, or confounding factors associated with both the exposures and outcomes. The randomized controlled trial (RCT) is considered to be the gold standard approach to test for causality. For instance, LDL cholesterol (LDL-c) was initially found to be associated with coronary artery disease (CAD) in an observational study ^5^, and the association was subsequently confirmed to be causal by RCTs ^6,7^. However, RCTs are time-consuming, expensive, and sometimes impractical or even unethical ^8^. It is not feasible to design RCTs that can test many different interventions simultaneously. Mendelian Randomization (MR) is an instrumental variable analysis that uses genetic variants, which are expected to be independent of confounding factors, as instrumental variables to test for causality ^9-11^. MR can be used to infer credible causal associations when RCTs are not feasible or as a strategy to rank order candidate causal associations to be prioritized for follow-up in RCTs. MR is becoming increasingly efficient and cost-effective given the ever-growing data curated from recent genome-wide association studies (GWAS). The large amount of GWAS data available in the public domain provide a great opportunity for methods that are able to make inference about causality by integrating summary-level GWAS data from different studies ^12-16^. We have previously shown that the power of MR could be greatly improved by a flexible analysis of summary-level GWAS data for exposure (e.g. risk factor) and outcome (e.g. disease) from two samples of large sample size (summary-data-based MR, SMR), and applied the SMR method to test if the effects of genetic variants on a phenotype are mediated by gene expression ^17^.

In this study, we extend the SMR approach to a more general form (generalized SMR or GSMR) by leveraging power from multiple genetic variants accounting for linkage disequilibrium (LD) between the variants, and demonstrate by simulation that GSMR is more powerful than existing summary-data-based MR methods ^12,13^. Separation of signals of causality from pleiotropy (a single locus directly affecting multiple phenotypes, also called type-II pleiotropy ^18^) and further separation of marginal effect from conditional effect (the net effect of a risk factor on the outcome accounting for the effects of other risk factors, e.g. there is no effect of HDL cholesterol on CAD correcting for the other serum cholesterol levels ^19,20^) are recognized issues that require careful interpretation in MR analyses ^18^. We develop a method (HEIDI-outlier) to detect and eliminate genetic instruments that have apparent pleiotropic effects on both risk factor and disease, and another method (conditional GSMR) to estimate the effect of a risk factor on disease conditioning on the genetic values of other risk factors. All methods developed in this study only require summary-level data (with LD between genetic variants from a reference sample with individual-level data), providing a great flexibility to integrate data from multiple studies. We apply the methods to publicly available data of very large sample sizes (*n* = up to 405,072 for risk factors and 184,305 for diseases) to test for the causal associations between health risk factors such as BMI, serum cholesterol levels and blood pressure levels and a range of human common diseases.

## Results

### Overview of the methods

Let *y* = the liability of a disease on the logit scale, *x* = a risk factor in standard deviation (SD) units and *z* = the genotype of a SNP (coded as 0, 1 or 2). The MR estimate of the causal effect of risk factor on disease ^9^ is *b̂_xy_* = *b̂_zy_/b̂_zx_*, where *b_zy_* is the effect of *z* on *y* on the logit scale (logarithm of odds ratio, logOR), *b_zx_* is the effect of *z* on *x*, and *b_xy_* is the logOR of *x* on *y* free of confounding from non-genetic factors (see below for the caveats in interpreting an MR estimate). SMR is a flexible and powerful MR approach that is able to estimate and test the significance of *b_xy_* using the estimates of *b_zx_* and *b_zy_* from independent samples ^17^. If there are multiple independent (or nearly independent) SNPs associated with *x* and the effect of *x* on *y* is causal, then all the *x*-associated SNPs will have an effect on *y* through *x* (**Fig. 1a**). In this case, *b_xy_* at any of the *x*-associated SNPs is expected to be identical in the absence of pleiotropy ^13,16,21^ because all the SNP effects on *y* are mediated by *x* (**Fig. 1b**). Therefore, increased statistical power can be achieved by integrating the estimates of *b_xy_* from all the *x*-associated SNPs using a generalized least squares (GLS) approach (**Online Methods**). The GSMR method essentially implements SMR analysis for each SNP instrument individually, and then integrates the *b_xy_* estimates of all the SNP instruments by GLS, accounting for sampling variance in *b̂_zx_* and *b̂_zy_* for each SNP and LD among SNPs. We demonstrate using simulations that there is no inflation in the GSMR test-statistics under the null hypothesis that *b_xy_* = 0 (**Supplementary Fig. 1**), that the estimate of *b_xy_* from GSMR is unbiased under the alternative hypothesis that *b_xy_* ≠ 0 (**Supplementary Table 1**), and that *b_xy_* approximately equals to logOR (where OR is the effect of risk factor on disease in observational study without confounding) (**Supplementary Fig. 2**). In comparison with the existing summary-data-based MR methods ^12^, GSMR is more powerful as demonstrated by simulation (**Supplementary Fig. 3**) especially when the number of SNP instruments is large, and has the advantage of accounting for LD if the SNP instruments are not fully independent.

**Figure 1.**
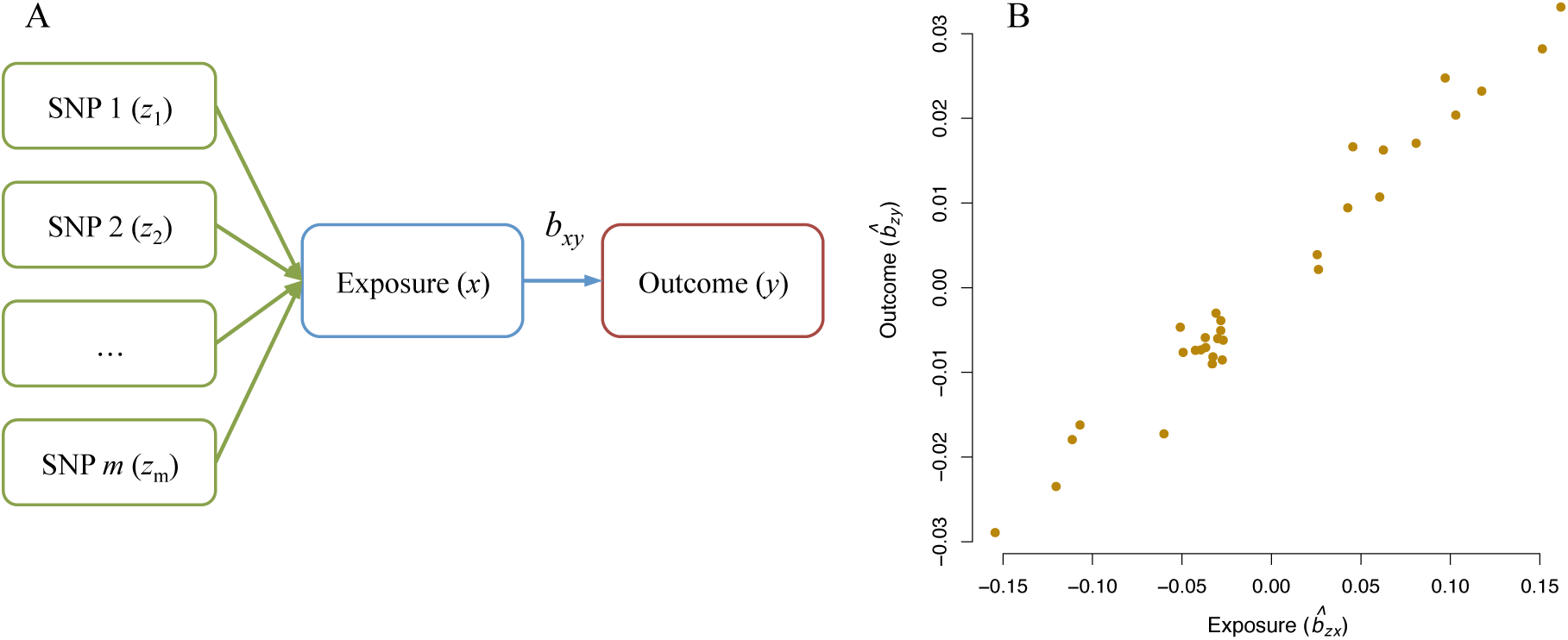
Leveraging multiple independent genetic instruments (*z*) to test for causality. Shown in panel (a) is a schematic example that if an exposure (*x*) has an effect on an outcome (*y*), any instruments (SNPs) causally associated with *x* will have an effect on *y*, and the effect of *x* on *y b_xy_*) at any of the SNPs is expected to be identical. This is further illustrated in a toy example in panel (b) that under a causal model, for the SNPs associated with *x*, the estimated effect of *z* on *y* (*b̂_zy_*) should be linearly proportional to the estimated effect of *z* on *x* (*b̂_zx_*) and the ratio between the two is an estimate of the mediation effect on *x* on *y*, i.e. *b̂_xy_* = *b̂_zy_/b̂_zx_*.

Pleiotropy is an important potential confounding factor that could bias the estimate and often leads to an inflated test-statistic in a MR analysis ^9,10,13,18^. We propose a method (called HEIDI-outlier) to detect the SNP outliers that are more consistent with pleiotropic effects on both exposure and outcome, and remove them from the GSMR analysis (**Online Methods** and **Supplementary Fig. 4**). We further develop an approximate method that only requires summary data to conduct a conditional GWAS analysis for a phenotype given other phenotypes. The purpose of developing this method is to estimate the effect of a risk factor on disease correcting for other risk factors (**Online Methods** and **Supplementary Fig. 5**), which helps to infer whether the marginal effect of the risk factor on disease depends on other risk factors or not, and to predict the joint effect of multiple risk factors on disease.

### Causal associations between 7 health risk factors and common diseases

We applied the methods to test for causal associations between 7 health risk factors and common diseases using data from multiple large studies. The risk factors are BMI, waist-to-hip ratio adjusted for BMI (WHRadjBMI), HDL cholesterol (HDL-c), LDL-c, triglycerides (TG), systolic blood pressure (SBP) and diastolic blood pressure (DBP). We chose these risk factors because of the availability of summary-level GWAS data from large samples (*n* = 108,039 to 322,154) (**Supplementary Table 2**). We accessed data for BMI, WHRadjBMI, HDL-c, LDL-c and TG from published GWAS ^22-24^ and data for SBP and DBP from the subgroup of UK Biobank (UKB) ^25^ with genotyped data released in 2015. We selected near-independent SNPs at a genome-wide significance level (*P_GWAS_* < 5 × 10^−8^) using the clumping algorithm (*r*^2^ threshold = 0.05 and window size = 1 Mb) implemented in PLINK ^26^ (**Online Methods**). Note that the GSMR method accounts for the remaining LD not removed by the clumping analysis. There were *m* = 84, 43, 159, 141, 101, 28 and 29 near-independent SNPs for BMI, WHRadjBMI, HDL-c, LDL-c, TG, SBP and DBP, respectively, after clumping. The summary-level GWAS data for the diseases were computed from two independent community-based studies with individual-level SNP genotypes, i.e. the Genetic Epidemiology Research on Adult Health and Aging ^27^ (GERA) (*n* = 53,991) and the subgroup of UKB ^25^ (*n* = 108,039). We included in the analysis 22 common diseases as defined in the GERA data (**Supplementary Table 3**). We added an additional phenotype related to comorbidity by counting the number of diseases affecting each individual, i.e. Disease Count, as a crude index to measure the general health status of an individual (**Supplementary Table 3**). We performed genome-wide association analyses of the 23 disease phenotypes in GERA and UKB separately (**Online Methods**). We assessed the genetic heterogeneity of a disease between the two cohorts by a genetic correlation (*r_g_*) analysis using the bivariate LD score regression (LDSC) approach ^28^. The estimates of *r_g_* across all the diseases (excluding Disease Count) varied from 0.75 to 0.99 with a mean of 0.91 (**Supplementary Table 3**), suggesting strong genetic overlaps for the diseases between the two cohorts. We therefore meta-analyzed the data of the two cohorts to maximize power using the inverse-variance meta-analysis approach ^29^.

Since the GSMR method only requires summary-level data, we further applied the GSMR analysis to 11 diseases for which there were summary data available from published case-control studies (*n* = 18,759 to 184,305) (**Supplementary Table 4**). The estimated SNP effects and SE for age-related macular degeneration (AMD) were not available in the summary data ^30^, which were estimated from z-statistics using an approximate approach (**Supplementary Note**). We applied the HEIDI-outlier approach to remove SNPs that showed pleiotropic effects on both risk factor and disease, significantly deviated from a causal model (**Online Methods**). Although this method might not be fail-safe against all threats to causality ^18,31^ and could potentially reduce power of GSMR (we use a threshold p-value of 0.01 for the HEIDI-outlier analysis which removes 1% of the SNPs by chance if there is no pleiotropic outlier) (**Supplementary Fig. 4**), it enhances the credibility of the analysis. The LD correlations between pairwise SNPs were estimated from the Atherosclerosis Risk in Communities (ARIC) data ^32^ (*n* = 7,703 unrelated individuals) imputed to 1000 Genomes (1000G) ^33^. Using the large data sets, we identified from GSMR analyses 45 significant causative associations between risk factors and diseases (**Supplementary Table 5** and **Fig. 2**). We controlled the family-wise error rate (FWER) at 0.05 by Bonferroni correction for 231 tests (*P_GSMR_* threshold = 2.16 × 10^-4^).

**Figure 2.**
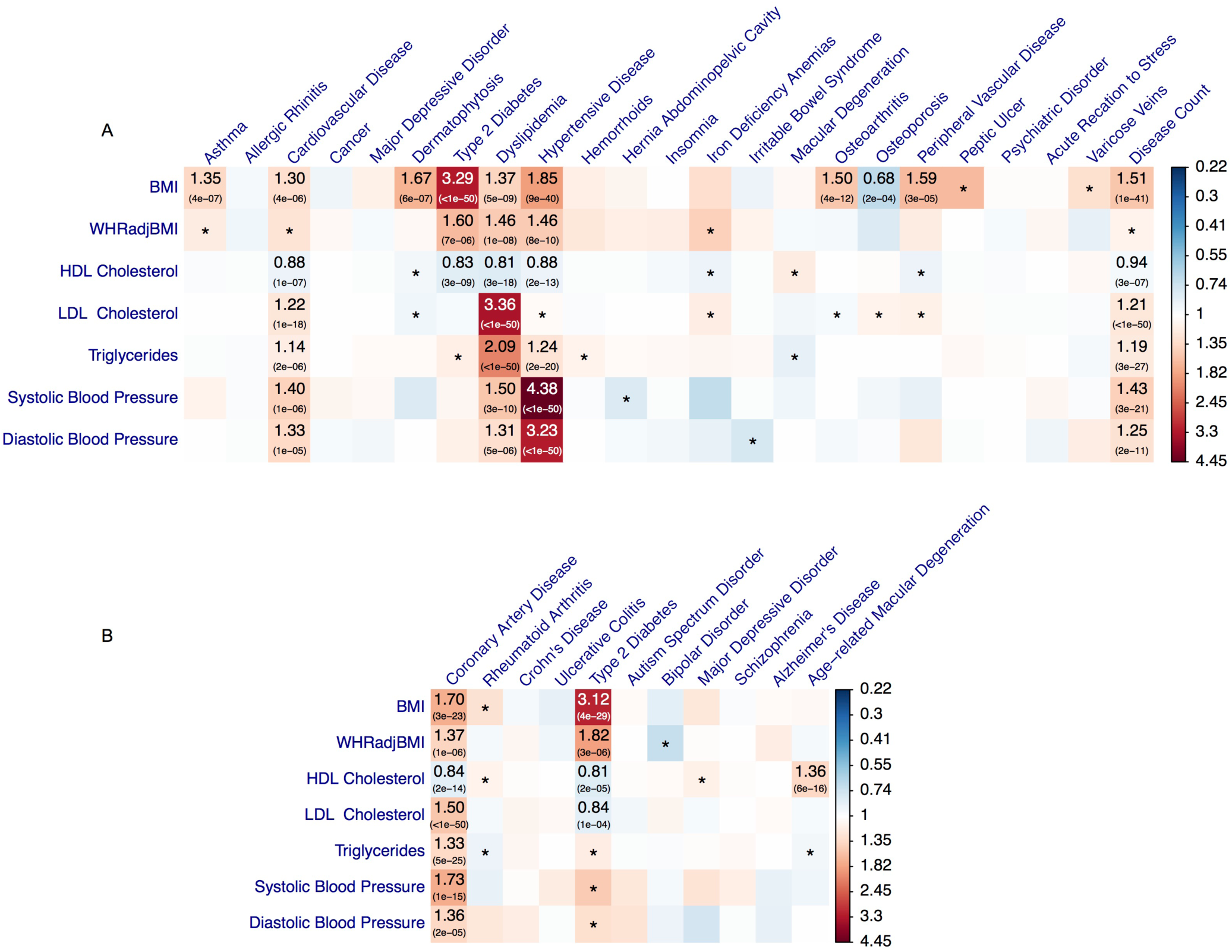
The association between genetic instruments that predict 7 modifiable risk factors and selected common diseases. Shown are the results from GSMR analyses with disease data (a) from a meta-analysis of two community-based studies (GERA and UKB) and (b) from published independent case-control studies (**Supplementary Table 3**). Colors represent the effect sizes (as measured by odds ratios, ORs) of risk factors on diseases, red for risk effects and blue for protective effects. The significant effects after correcting for multiple testing (*P*_GSMR_ < 2.2 × 10^−4^) are labeled with ORs (p-values). The nominally significant effects (*P*_GSMR_ < 0.05) are labeled with “∗”

### Obesity and common diseases

Results from analyses of the community-based data showed that BMI had risk effects on T2D (odds ratio, OR = 3.29), hypertensive disease (OR = 1.85), dermatophytosis (i.e. tinea) (OR = 1.67), peripheral vascular diseases (PVD) (OR = 1.59), osteoarthritis (OR = 1.50), dyslipidemia (OR = 1.37), asthma (OR = 1.35) and CVD (OR = 1.30). The risk effects of BMI on T2D, CVD and hypertensive disease have been confirmed by RCT ^34^ (**Supplementary Table 5**), providing proof-of-principle validation. The interpretation of OR_(BMI→T2D)_ = 3.29 is that people whose BMI are 1 SD (SD = 3.98 for BMI in European men corresponding to ~12kg of weight for men of 175cm stature; see **Supplementary Table 6** for SD for the other risk factors) above the population mean will have 3.29 times increase in risk to T2D compared with the population prevalence (~8% in the US). It is interesting to note that the estimate of *b_xy_* at the *TCF7L2* locus strongly deviated from those at the other loci (**Fig. 3**), suggesting that the *TCF7L2* SNP has pleotropic effects on BMI and T2D. The *TCF7L2* SNP was detected as an outlier by the HEIDI-outlier method and removed from the GSMR analysis. In addition, the risk effect of BMI on asthma is in line with the result from a recent MR study (using a weighted genetic allele score as the instrument) that higher BMI increases the risk of childhood asthma ^35^. Moreover, we identified a protective effect of BMI against osteoporosis (OR = 0.68), consistent with the observed associations in previous studies ^36,37^.

**Figure 3.**
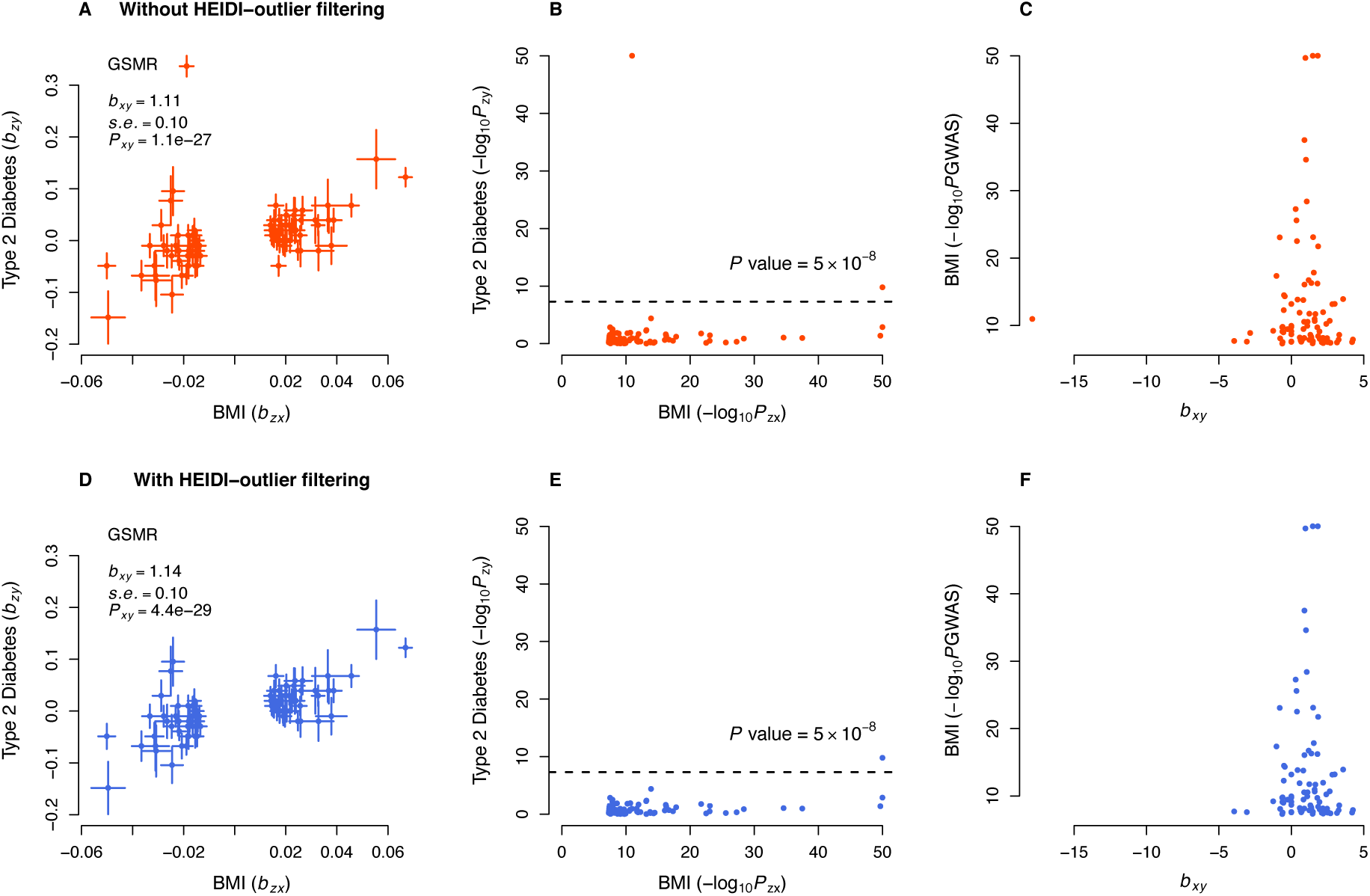
GSMR analysis to test for the effect of BMI on T2D with and without filtering the pleiotropic outliers. Shown in panels (a) and (b) are the plots of effect sizes and association p-values of all the genetic instruments from GWAS for BMI vs. those for T2D. Shown in panel (c) is the plot of *b_xy_* vs. GWAS p-value of BMI at each genetic variant. Shown in panels (d), (e) and (f) are the plots for the instruments after the pleiotropic outliers being removed by the HEIDI-outlier approach (see **Online Methods** for details of the HEIDI-outlier approach). The dashed lines in panels (b) and (e) represent the GWAS threshold p-value of 5 × 10^−8^. The coordinates in panels (b), (c), (e) and (f) are truncated at 50 for better graphic presentation.

The estimated risk effect of BMI on T2D in the community data (OR = 3.29) was similar to that in the case-control data (OR = 3.12, **Fig. 2b** and **Supplementary Table 5**). We also observed a strong risk effect of BMI on coronary artery disease (CAD) in the case-control data (OR = 1.70), in line with the risk effect of BMI on CVD (OR = 1.30) in the community data. Our result confirms a negligibly small effect of BMI on major depressive disorder (MDD), consistent with a previous MR study that uses a SNP-derived genetic risk score as a single instrument in a smaller sample ^38^, while noting that the MDD GWAS data are currently underpowered as assessed by the limited number of genome-wide significant (GWS) loci currently detected.

Being overweight is a risk factor for general health outcomes as indicated by its risk effect on Disease Count (*b̂_xy_* = 0.41) in the community data. The question is then how *b_xy_* for Disease Count should be interpreted. We have shown in **Supplementary Fig. 6** that the estimate of *b_xy_* for Disease Status (a dichotomous phenotype to indicate whether an individual is affected by any of the 22 diseases) was very similar to that for Disease Count. Although Disease Status and Disease Count are two distinct phenotypes and the analysis of Disease Count is more powerful, for the ease of interpretation, *b_xy_* for Disease Count can be approximately interpreted as logOR for Disease Status. Hence, *b̂_xy_ =* 0.41 for Disease Count is approximately equivalent to OR = 1.51 for Disease Status, meaning an increase of BMI by 1 SD will increase the probability of being affected by any of the 22 diseases by a factor of ~1.5.

We included in the analysis two obesity-related traits, BMI and WHRadjBMI. BMI is a measure of the amount of tissue mass and WHRadjBMI is a measure of body fat distribution. Previous studies suggest that BMI- and WHRadjBMI-associated genetic loci are enriched for genes expressed in central nervous system ^22^ and adipose tissue ^23^, respectively, which implies potentially different genetic aetiologies of the two traits. We found that the effects of WHRadjBMI and BMI on disease were largely concordant (**Supplementary Fig. 7a**), suggesting that the genetic etiologies of the two traits may differ but their effects on health outcomes are similar. Note that WHRadjBMI has been adjusted for BMI so that that effect sizes of WHRadjBMI on diseases should be independent of BMI. Nevertheless, the effect sizes of WHRadjBMI on diseases were smaller than those of BMI, and WHRadjBMI was detected with significant effects on only 4 diseases (T2D, dyslipidemia, hypertension and CAD) (**Fig. 2**), although the smaller number of detections could be because of the smaller number of instruments used for WHRadjBMI (*m* = 43) than for BMI (*m* = 84) as the power of GSMR is proportional to the number of instruments (**Supplementary Fig. 3**).

### “Good” vs. “bad” cholesterol

LDL-c is a known causative risk factor for CAD as confirmed by RCTs ^6,7^. We found that LDL-c had a significant risk effect on dyslipidemia (OR = 3.36) and CVD (OR = 1.22) in the community data, and CAD (OR = 1.50) in the case-control data (**Fig. 2**). TG had a significant risk effect on dyslipidemia (OR = 2.09), hypertensive disease (OR = 1.24) and CVD (OR = 1.14) in the community data, and CAD (OR = 1.33) in the case-control data (**Fig. 2**). The effects of TG on diseases were largely consistent with those for LDL-c (**Supplementary Fig. 7b**), despite the modest phenotypic correlation between the two traits (*r_p_* = 0.19 in the ARIC data). Both LDL and TG had significant risk effects on Disease Count in the community data (**Fig. 2**).

There was another example where the HEIDI-outlier approach detected strong effects due to pleiotropy. The effect of LDL-c on Alzheimer’s disease (AD) was highly significant without HEIDI-outlier filtering (OR = 1.35 and *P_GSMR_* = 7.8 × 10^−16^) (**Fig. 4**). The HEIDI-outlier analysis flagged 16 SNPs, 12 of which are located in the *APOE* gene region (LD *r*^2^ among these SNPs < 0.05) and all of which had highly significant effects on both LDL-c and AD. Excluding these SNPs makes a more conservative GSMR test because if there is a true causal relationship of increased LDL-c with AD, then the GSMR test should remain significant based on evidence from other LDL-c associated SNPs. In fact, after removing the 16 pleiotropic SNPs, the effect of LDL-c on AD was not significant (OR =1.03, *P_GSMR_* = 0.47). Nevertheless, the multiple pleiotropic signals clustered at the *APOE* locus are worth further investigation (**Supplementary Fig. 8**).

**Figure 4.**
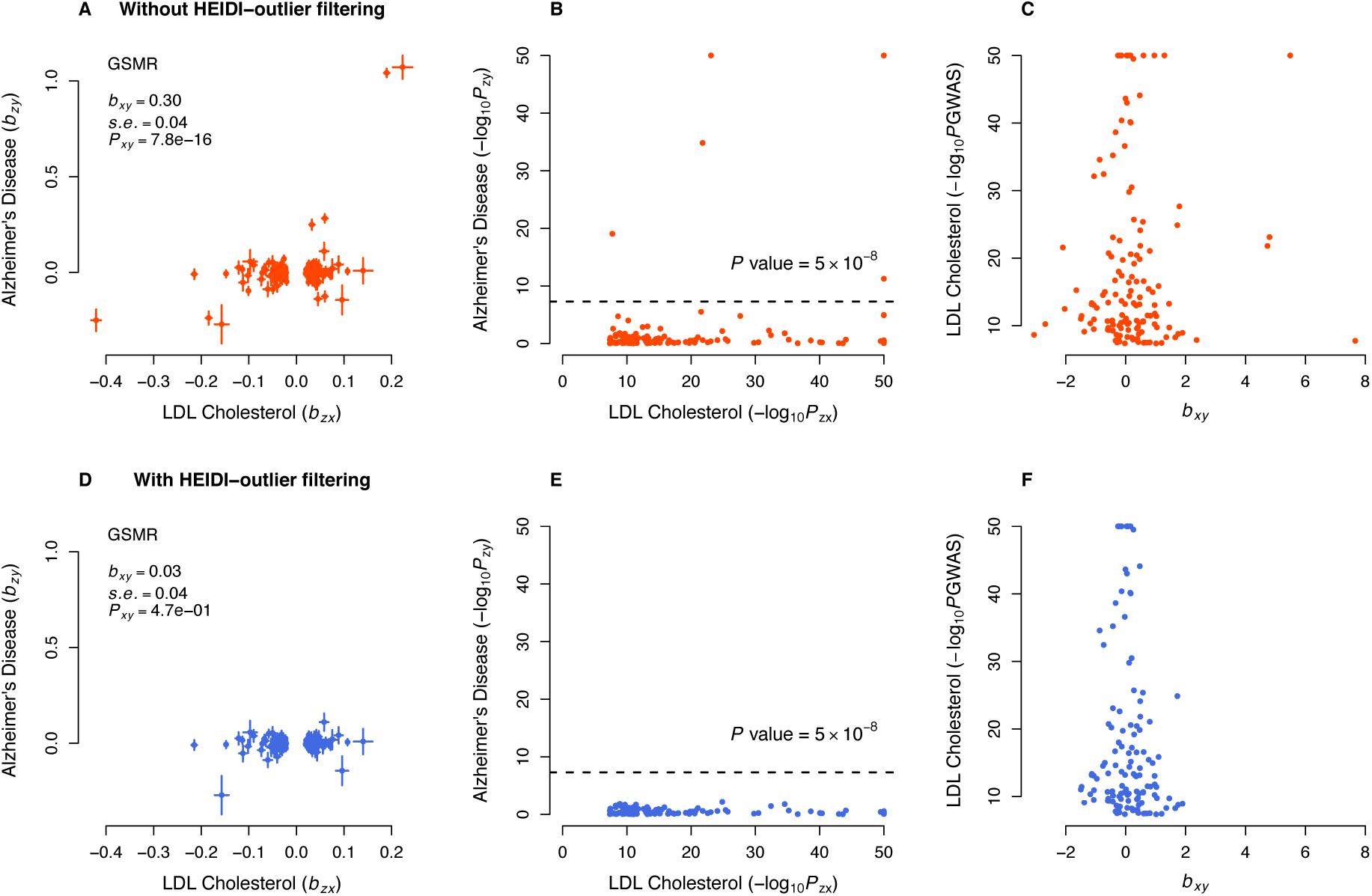
GSMR analysis to test for the effect of LDL-c on Alzheimer’s disease (AD) with and without pleiotropic outliers. Shown in panels (a) and (b) are the plots of effect sizes and association p-values of the original set of instruments from GWAS for LDL-c vs. those for AD. Shown in panel (c) is the plot of *b_xy_* vs. GWAS p-value of LDL-c at each genetic variant. Shown in panels (d), (e) and (f) are the plots for the instruments after the pleiotropic outliers being removed by the HEIDI-outlier approach (see **Online Methods** for details of the HEIDI-outlier approach). The dashed lines in panels (b) and (e) represent the GWAS threshold p-value of 5 × 10^−8^. The coordinates in panels (b), (c), (e) and (f) are truncated at 50 for better graphic presentation.

We identified a significant protective effect of LDL-c against T2D (OR = 0.84, *P_GSMR_* = 1.1 × 10^−4^) in the case-control data, which might explain the observation from a previous study that lowering LDL-c using statin therapy is associated with a slightly increased risk of T2D ^39^. The estimate was not significant in the community data (likely due to the lack of power) but in a consistent direction (OR = 0.95, *P_GSMR_* = 0.08). Given the strong genetic correlation between the two T2D data sets (*r_g_* = 0.98, SE = 0.062) as estimated from the bivariate LDSC analysis ^28^, we meta-analyzed the two data sets using the inverse-variance approach, and performed the GSMR analysis to re-estimate the effect of LDL-c on T2D using the T2D meta-analysis data. The effect size was highly significant (OR = 0.88, *P*_GSMR_ = 3.0 × 10^-7^).

The consequences of HDL-c on health outcomes are controversial ^40^. Observational studies suggest that HDL-c is associated with a reduced risk to CAD ^41^ whereas genetic studies show that the effect of HDL-c on CAD is not significant conditional on LDL-c and TG ^19,20^. We found that HDL-c had protective effects against T2D (OR = 0.83), hypertensive disease (OR = 0.88), CVD (OR = 0.88) and Disease Count (OR = 0.94) in the community data, and T2D (OR = 0.81) and CAD (OR = 0.84) in the case-control data. However, none of these effects remained significant conditioning on the other risk factors, suggesting that the marginal effects of HDL-c on diseases are dependent of the other risk factors (see below for details of the results from conditional analyses). The effect of HDL-c on dyslipidemia is negative (*b̂_xy_* = –0.21 and OR = 0.81), which is obvious because one of the diagnostic criteria for dyslipidemia is an abnormally low level of HDL-c. In addition, there was a highly significant risk effect (OR = 1.36) of HDL-c on age-related macular degeneration (AMD) in the case-control data. The associations between lipids and AMD are controversial and results from different observational studies are inconsistent ^42^. Our results support the observations that increased HDL-c is associated with increased risk of AMD ^42-44^. It should also be noted that LDL-c and TG also appeared to be associated with AMD before HEIDI-outlier filtering but the effects were not significant after HEIDI-outlier filtering (**Supplementary Fig. 9**), implying that the observed association between LDL-c (or TG) and AMD in epidemiological studies ^42^ might be due to pleiotropy.

### Blood pressure and common diseases

We identified significant risk effects of SBP on hypertensive disease (OR = 4.38), dyslipidemia (OR = 1.50), CVD (OR = 1.40) and Disease Count (OR = 1.43) in the community data, and CAD (OR = 1.73) in the case-control data. The results for SBP and DBP were highly concordant (**Fig. 2** and **Supplementary Fig. 7c**). The risk effect of blood pressure on CAD is known to be causal as confirmed by RCTs ^45,46^. Note that the power of the GSMR analysis for blood pressure was likely to be limited given the small number of instruments used (*m* < 30).

### Conditional effects of risk factors on diseases

We have identified (from the analyses above) 45 significant causal associations between health risk factors and diseases (**Fig. 2**). Since the risk factors are not independent, we further sought to estimate the effect of a risk factor on a disease conditioning on other risk factors, which helps to infer the mediating effects between risk factors ^18^. To do this, we first performed GSMR analysis to test for causal associations among the risk factors. We detected 19 significant associations among the 7 risk factors at a FWER of 0.05 (*P*_GSMR_ < 1.2 × 10^−3^) (**Supplementary Fig. 10**). For example, BMI had a significant negative effect on HDL-c (*b̂_xy_* = –0.29), and positive effects on TG (*b̂_xy_* = 0.28) and DBP (*b̂_xy_* = 0.15).

We developed a novel approach called mtCOJO (<u>m</u>ulti-<u>t</u>rait-based <u>co</u>nditional and <u>jo</u>int analysis) to perform a GWAS analysis for a trait conditioning on other traits using GWAS summary data (**Online Methods** and **Supplementary Fig. 5**). We then re-ran the GSMR analysis using GWAS summary data from the mtCOJO analysis (**Online Methods**). The mtCOJO analysis requires the estimates of *r_g_* among the covariate risk factors, *r_g_* between covariate risk factors and disease, SNP-based heritability (
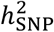)
for covariate risk factors and disease, and effect sizes of covariate risk factors on the target risk factor and disease, all of which can be computed from summary data using the univariate ^47^ and bivariate ^28^ LDSC approaches (**Online Methods**, and **Supplementary Tables 8**-**10**). Given the similar GSMR results between BMI and WHRadjBMI and between SBP and DBP (**Supplementary Fig. 7**), we did not include DBP and WHRadjBMI in the conditional analysis to avoid over-correction.

Results from conditional analyses were largely consistent with those from unconditional analyses (**Fig. 5** and **Supplementary Table 11**), suggesting that most of the marginal effects are independent of the other risk factors analyzed in this study. Conditioning on the other risk factors, SBP, LDL-c and BMI were the three major risk factors for CAD, BMI was still a large risk factor for T2D and the protective effect of LDL-c on T2D remained largely unchanged (**Supplementary Fig. 11**). The effect of BMI on T2D and the effect of TG on dyslipidemia decreased substantially in the conditional GSMR analyses (**Supplementary Table 11**), suggesting that these effects partly depend on the other risk factors.

**Figure 5.**
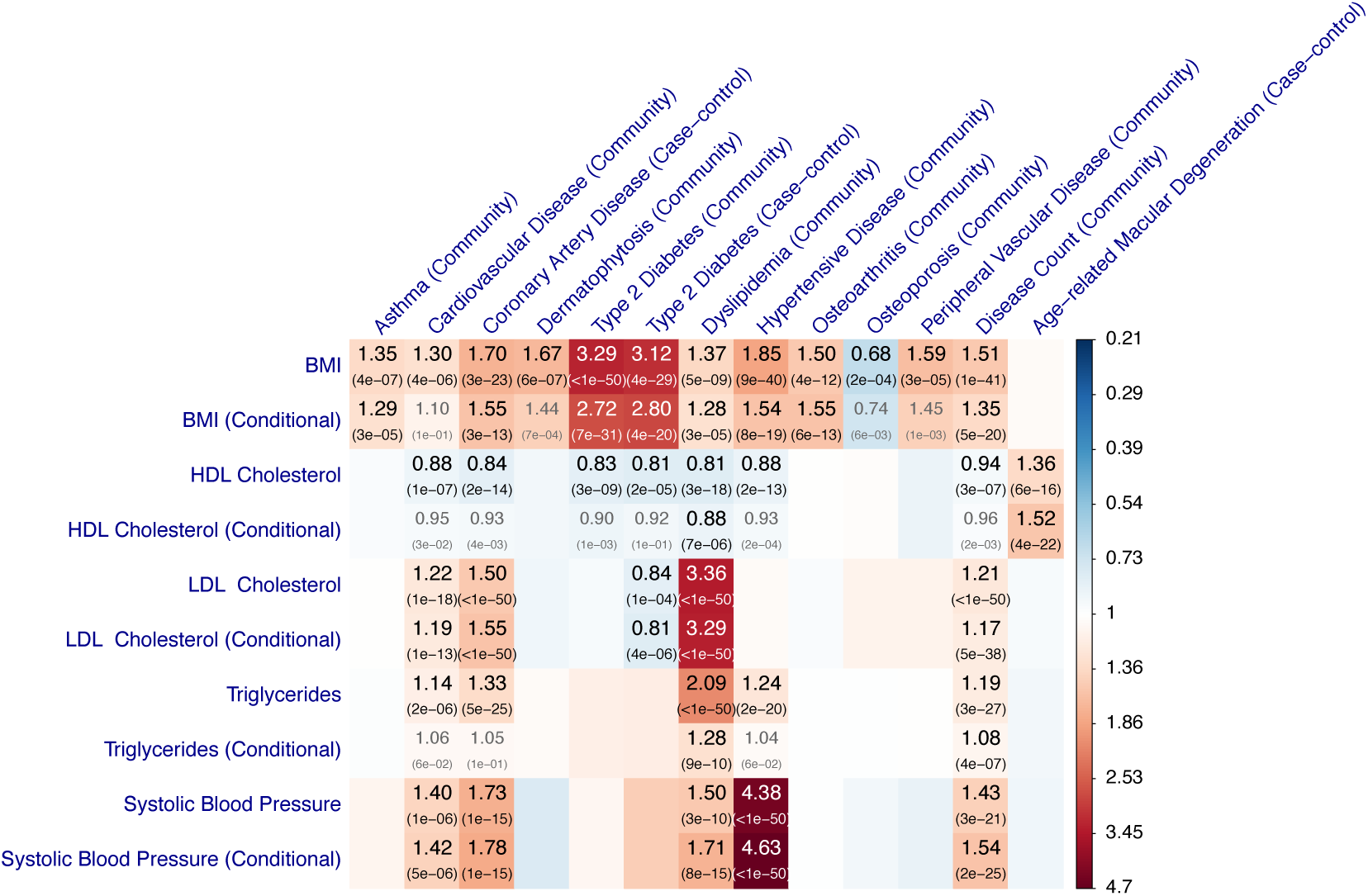
GSMR vs. conditional GSMR. Shown are the results from the GSMR analyses compared with those from the conditional GSMR analyses. In the conditional GSMR analysis, the effect size of each risk factor on disease was estimated conditioning on the other risk factors (see **Online Methods** for details of the conditional method). “Community”: disease GWAS data from a meta-analysis of the two community-based studies. “Case-control”: disease GWAS data from independent published case-control studies. In gray are the associations that do not pass the p-value threshold 2.16 × 10^−4^ in the conditional analysis.

We show above that the GSMR analyses identified significant protective effects of HDL-c against CVD, CAD, T2D and hypertension (**Supplementary Fig. 12**). However, all the effects became non-significant conditional on the covariate risk factors (i.e. BMI, LDL-c, TG and SBP), suggesting that the marginal effects of HDL-c on the diseases were likely to be mediated or driven by the covariates due to the complex bidirectional causative associations between HDL-c and the other risk factors as illustrated in **Supplementary Fig. 10**. It is difficult to distinguish whether the effects of HDL-c on the diseases are mediated or driven by the covariates (**Supplementary Fig. 13**) because both HDL-c and some of the covariates showed a significant effect on the diseases in either unconditional or conditional GSMR analysis (**Fig. 5**). Nevertheless, there might be an exception, that is, the association between HDL-c on AMD, because HDL-c is the only risk that showed a significant effect on AMD (OR = 1.36 with *P*_GSMR_ = 5.9 × 10^−16^) and the effect size remained highly significant conditioning on the covariates (conditional OR = 1.52 with *P*_GSMR_ = 4.4 × 10^−22^). We hypothesize that HDL-c is likely to be a direct risk factor for AMD and the effect size is largely independent of the covariate risk factors analyzed in this study. Note that the estimate of conditional effect was slightly larger than that of marginal effect, possibly because part of the marginal effect of HDL-c on AMD was masked by the negative effects of the covariates on HDL-c (**Supplementary Fig. 10**).

Given the estimates from conditional GSMR analyses (**Fig. 5 and Supplementary Table 11**), we could use an approximate approach to calculate the aggregate effect of multiple risk factors on a disease, i.e. log(OR = ∑[*x_i_* log(OR_*i*_)]. Here is a hypothetical example. If all the risk factors increase by 1 SD (i.e., ~4 kg/m^2^ for BMI, ~1 mmol/L for LDL-c, ~1 mmol/L for TG and ~19 mmHg for SBP), we would have an increased risk of approximately 2.3-fold to T2D (e^1.03-0.21^), and 4.3-fold to CAD (e^0.44+0.44+0.58^).

### Effects of other phenotypes on diseases

Having identified a number of causal associations between 7 modifiable risk factors and common diseases, we then sought to test whether there were causative associations between other phenotypes and diseases. We included in the analysis two traits, height ^48^ and years of schooling ^49^ (EduYears), for which there were a large number of instruments owing to the large GWAS sample sizes. We selected 811 and 119 near-independent GWS SNPs for height and EduYears, respectively, using the clumping analysis (**Online Methods**). The threshold *P*_GSMR_ after Bonferroni correction was 7.6 × 10^−4^ correcting for 66 tests. The large number of instruments for height gave us sufficient power to detect a small effect. Results showed that height had significant protective effects against irritable bowel syndrome (OR = 0.82), dyslipidemia (OR = 0.84), osteoporosis (OR = 0.85), acute reaction to stress (OR = 0.85), hypertensive disease (OR = 0.86), T2D (OR = 0.86) and asthma (OR = 0.90) in the community data, and CAD (OR = 0.77), T2D (OR = 0.84), AD (OR = 0.85) and AMD (OR=0.91) in the case-control data (**Fig. 6** and **Supplementary Table 12**). It is interesting to note that height was protective against AD which was not affected by any of the 7 health risk factors described above. Results also showed that height had a risk effect on varicose veins (OR = 1.38), dermatomycosis (OR = 1.24), peripheral vascular disease (PVD, OR = 1.15), cancer (OR = 1.09), osteoarthritis (OR = 1.09) and CVD (OR = 1.07) in the community data, and rheumatoid arthritis (RA, OR = 1.29) in the case-control data. The inconsistent effects of height on CAD (OR = 0.77) and CVD (OR = 1.07) implies heterogeneous effects of height on vascular outcomes consistent with results from some of the observational studies ^50-52^ and a previous MR study ^53^. Height was the only risk factor that showed a significant risk effect on cancer (*P*_GSMR_ = 3.1 × 10^−7^) in this study, in line with the results from previous observational and MR studies ^54,55^.

**Figure 6.**
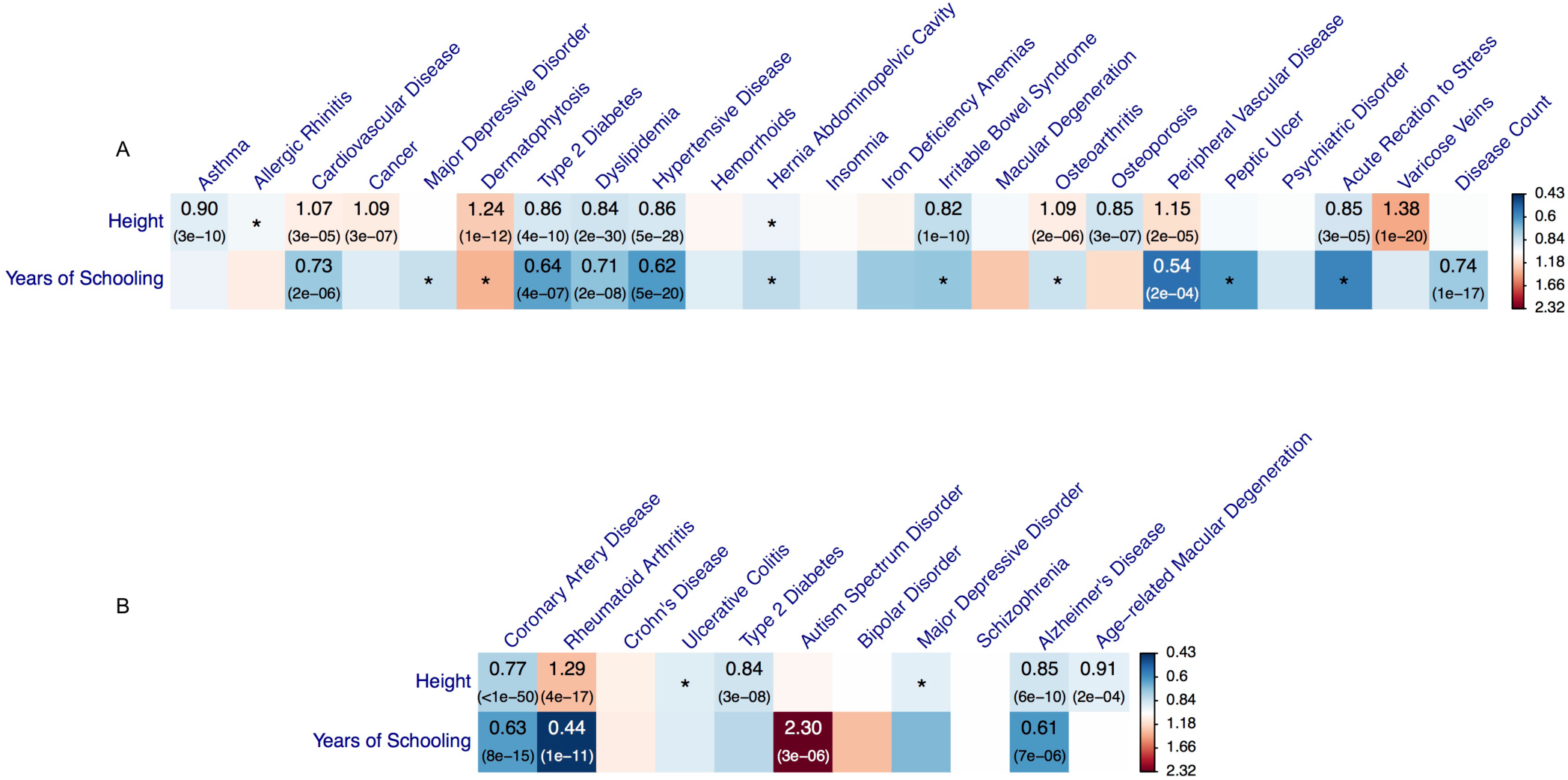
Effects of height and educational attainment on common diseases. Shown are the results from GSMR analyses with disease data (a) from a meta-analysis of the GERA and UKB studies and (b) from published independent case-control studies. Colors represent the effect sizes (as measured by odds ratios, ORs) of risk factors on diseases, red for risk effects and blue for protective effects. The significant effects after correcting for multiple testing (*P*_GSMR_ < 7.6 × 10^−4^) are labeled with ORs (p-values). The nominally significant effects (*P*_GSMR_ < 0.05) are labeled with “∗”.

Our results also showed that EduYears had protective effects against almost all the diseases (**Fig. 6** and **Supplementary Table 12**). It showed protective effect against PVD (OR = 0.54), hypertensive diseases (OR = 0.62), T2D (OR = 0.64), dyslipidemia (OR = 0.71) and CVD (OR = 0.73) in the community data, and RA (OR = 0.44), AD (OR = 0.61) and CAD (OR = 0.63) in the case-control data. It also showed significant protective effect on Disease Count (OR = 0.74), suggesting that educational attainment is protective for general health outcomes. The protective effect of EduYears against AD is consistent with the observed association from epidemiological studies ^56^. On the other hand, however, EduYears showed a strong risk effect on autism spectrum disorder (ASD, OR = 2.30), which is not influenced by SNP outliers (**Supplementary Fig. 14**) and consistent with a positive estimate of genetic correlation (*r_g_* = 0.28, SE = 0.038) from a bivariate LD score regression analysis. Since ASD is mostly diagnosed in childhood prior to completion of education, the risk effect of EduYears on ASD is probably because EduYears is a genetic proxy for IQ. While ASD is associated with cognitive deficits in executive function, the relationship with IQ is complex ^57^. Our results are consistent with those from a Danish study of more than 160,000 male conscripts ^58^ which found that brothers of those with ASD had a significantly *higher* than average IQ score (whereas brothers of those with every other recorded psychiatric disorder had significantly lower than average IQ scores). We note that GSMR analysis tests the effect of EduYears on ASD rather the reverse direction. It is possible that the effects are bi-directional and opposite (see below for such examples from the reverse GSMR analysis), suggesting a U-shaped relationship between ASD and cognition.

### Reverse GSMR analysis

It is important to note that the causative associations identified from the GSMR analyses above are unlikely to be explained by reverse causality for two reasons. First, the individuals used in GWAS for risk factors were independent of the individuals used in GWAS for diseases (the only exception was that the blood pressure GWAS data set was part of the community-based disease GWAS data). Secondly, if the associations presented above are driven by reverse causality, we would expect to see strong association signals of the instruments with the diseases, which is not the case as demonstrated in **Supplementary Fig. 15**, an idea not too dissimilar to the asymmetry analysis ^21^ that has been used to infer causality in a previous study ^16^. Nevertheless, it is interesting to investigate the changes in risk factors after development of the diseases. To do this, we selected instruments for diseases from the disease GWAS data (note that the instruments used in the reverse-GSMR analysis were distinct from those used in the forward-GSMR analysis). We performed a reverse-GSMR analysis of the risk factors and diseases for which there was a significant association in the forward-GSMR analysis above (**Supplementary Note**). We identified 10 significant reverse effects (i.e. the effect of disease on risk factor) in the community data and 4 in the case-control data at a FWER of 0.05 (P_reverse-GSMR_ < 1.0 × 10^−3^) (**Supplementary Table 13**). The estimates of reverse effects were very small compared with those of the forward effects. Interestingly, there were two cases where the estimated forward and reverse effects were in opposite directions, i.e. *b̂*_*xy*(BMI→T2D)_ = 1.19 and *b̂*_*xy*(T2→BMI)_ = – 0.07 (*P* = 3.6 × 10^−26^); *b̂*_*xy*(BMI→dyslipidemia)_ = 0.32 and *b̂*_*xy*(dyslipidemia→BMI)_ = –0.03 (*P* = 2.0 × 10^−10^), meaning that although BMI is risk factor for the two diseases, patients who have developed the diseases tend to lose weight.

## Discussion

We proposed a flexible and powerful approach that performs a MR analysis with multiple near-independent instruments (i.e., GWS SNPs) to test for causal association between a risk factor (or phenotype) with a disease using summary-level GWAS data from independent studies. We have implemented the method in an R package (**URLs**). The method and software tool are general and can be applied more broadly to test for causality in other fields such as behavioral sciences. We applied the method to summary data from GWAS of very large sample size, and identified a large number of causal associations between risk factors and common diseases. As the effect sizes of SNPs on risk factor and disease used in the GSMR analysis were from independent GWAS data sets, the effect of risk factor on disease estimated from GSMR was very unlikely to be confounded by environmental factors. The result, however, could be biased if there are SNPs that have strong pleiotropic effects on both risk factor and disease. For example, the result for LDL-c and Alzheimer’s disease could have been biased due to 16 pleiotropic SNPs (**Fig. 4**). There are four lines of evidence that our results are not driven by pleiotropy between risk factor and disease. First, as demonstrated in the example above, we have used the HEIDI-outlier approach that could effectively remove instruments with strong putative pleiotropic effects (**Figs. 3** and **4**). After the HEIDI-outlier filtering, the instruments selected for risk factors did not show strong associations with the diseases except for those highly related diseases and traits (e.g. lipids and dyslipidemia; blood pressures and hypertensive disease) (**Supplementary Fig. 15**). Note that the test-statistics decreased slightly after filtering SNPs by HEIDI-outlier (**Supplementary Fig. 4b**), indicating that the result from the analysis with HEIDI-outlier filtering is more conservative. Second, if the results were driven by pleiotropy, we would expect the estimates of *b_xy_* from reverse-GSMR comparable with those from GSMR, which is not what we observed (**Supplementary Table 13**). Third, the estimates of *b_xy_* were highly consistent with the slopes from Egger regression that are considered to be free of confounding from pleiotropy ^13^ (MR-Egger) (**Supplementary Fig. 16**). Note that we used GSMR for the main analyses because in comparison with MR-Egger and inverse-variance weighted method (MR-IVW, equivalent to MR-Egger without intercept) ^12^, GSMR gains power by taking the sampling variation of *b̂_zx_* and *b̂_zy_* into account as demonstrated in simulations (**Supplementary Fig. 3**), and GSMR also has the advantage of accounting for LD among SNPs not removed by the clumping analysis, a property that is important specially when the number of instruments is large. Fourth, the intercepts from MR-Egger (a significant deviation of the intercept from 0 is evidence for the presence of pleiotropy) were very small relative to the slopes (**Supplementary Fig. 17**), and there was no inflation in the test-statistics (**Supplementary Figs. 17b and 17c**), suggesting that the degree of pleiotropy was negligible if there was any.

We have shown above that our results were not driven by pleiotropy and reverse-causality. In some cases, the relationship between a risk factor and a disease could be a mixture of multiple models. For example, we have shown above that BMI had a risk effect on T2D, which has been confirmed by RCT ^34^, that T2D had a significant reverse effect on BMI and effect size was negative, and that there was a SNP (at the *TCF7L2* gene locus) that appeared to have pleiotropic effects on T2D and BMI (**Fig. 3**), a mixture model of causality, reverse causality and pleiotropy. In addition, we demonstrated by the conditional GSMR analyses that the mediation effects (i.e. the effect size of a risk factor on disease mediated or driven by other risk factors) are apparently small for most risk factors except for HDL-c (**Fig. 5** and **Supplementary Table 11**).

Nevertheless, there are a few several caveats in interpreting the GSMR results. First, if the exposure is a composite trait that comprises multiple sub-phenotypes, we could not rule out the possibility that the effect of exposure on disease is driven by one of the sub-phenotypes. For instance, we have identified from the GSMR analysis that EduYears had effects on many diseases (**Fig. 6**). A conservative interpretation is that these are the effects of the genetic component of EduYears (e.g. cognitive ability and personality) on health outcomes. If we express EduYears = *g* + *e* where *g* is the genetic component of EduYears and *e* is the residual component that includes environmental influence, then the SNPs identified from GWAS for EduYears are those associated with *g* rather than *e*, meaning that the GSMR analysis for EduYears was performed on *g* rather than *e* and thus did not provide any evidence whether *e* also has effects on diseases. Therefore, strictly speaking, the causative associations identified in this study are not definitive and need to be confirmed by follow-up RCTs in the future, if practical. Second, the effect of a risk factor on disease can be non-linear (e.g. the relationship between BMI and mortality is a U-shaped curve ^3^, suggesting that both underweight and overweight are risk factors of death) whereas we used a linear approximation to estimate the effect because of the limited information that we had access to from GWAS summary data. Therefore, the *b_xy_* estimates need to be interpreted with cautions at extremes. Third, although we have identified a large number of associations, we would expect that associations of small effect size would be missed in our study (e.g. the instrument for SBP was based on only 28 SNPs). The power can be improved in the future with GWAS results based on larger sample sizes. Fourth, our analyses ignored age-specific and sex-specific effects because of the lack of data from age- and sex-stratified analyses. Last but not least, we used estimates of SNP effects estimated from population-based studies to approximate those in case-control studies (e.g. we used SNP effects from the GIANT meta-analysis to approximate those in the T2D case-control studies), which might explain the difference between the GSMR estimates in the community and case-control data (Fig. 2).

We present here summary-data-based MR analysis approaches that leverage the large amount of GWAS data from independent studies to detect the effect of a risk factor on diseases and assess the effect size conditional on the other risk factors. All the data used in this study were from the public domain, which demonstrates the power of an integrative analysis of existing data to make novel discoveries. The causal associations identified in this study not only provided important candidates to be prioritized in RCTs in the future but also provided fundamental knowledge to understand the biology of the diseases. Our findings of the effects of risk factors on common diseases could have a significant influence on medical research, pharmaceutical industry and public health.

## Online Methods

### The GSMR method

Mendelian randomization is a method that uses genetic variants as instrumental variables to test for causative association between an exposure and an outcome ^9^. Let *z* be a genetic variant (e.g. SNP), *x* be the exposure (e.g. health risk factor) and *y* be the outcome (e.g. disease). If *z* is significantly associated with *x*, the effect of *x* on *y* can be estimated using a two-step least squares (2SLS) approach ^59^

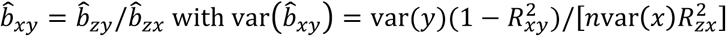

where *n* is the sample size,
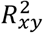
is the variance in *y* explained by *x*, and
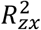
is the variance in *x* explained *z*. This analysis requires individual-level data so that the statistical power could be limited if *b_xy_* is small. We have previously proposed an approach that only requires summary-level data to estimate *b_xy_* so that the power can be greatly improved if *b_zy_* and *b_zx_* are estimated from independent studies of large sample size ^17^, i.e., *b̂_xy_* = *b̂_zy_/b̂_zx_* with
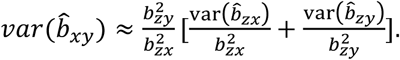
We called this approach a <u>s</u>ummary-data based <u>M</u>endelian <u>r</u>andomization (SMR) analysis ^17^. We have also shown previously that a SMR analysis using a single genetic variant is unable to distinguish between causality (the effect of SNP on outcome is mediated by exposure) and pleiotropy (the SNP has distinct effects on exposure and outcome). Here, we extend the SMR method to use all the top associated SNPs at a genome-wide significance level for the exposure as instrumental variables to test for causality. We call this method a generalized SMR (GSMR) analysis. The basic idea of GSMR is that if *x* is causal for *y*, any SNP associated with *x* will have an effect on *y*, and the expected value of *b̂*_*xy*(*i*)_ at any SNP *i* will be identical in the absence of pleiotropy. Let *m* be the number of GWS top SNPs associated with *x* after clumping. We have **b̂**_*xy*_ = {*b̂*_*xy*(1)_, *b̂*_*xy*(2)_, …, *b̂*_*xy*(*m*)_} with *b̂*_*xy*(*i*)_ = *b̂*_*zy*(*i*)_/*b̂*_*zx*(*i*)_, and **b̂**_*xy*_ ~ *N*(**1***b_xy_*, **V**) where **1** is an *m* × 1 vector of ones and **V** is the variance-covariance matrix of **b̂**_*xy*_. We have derived previously that the *ij*-th element of **V** is
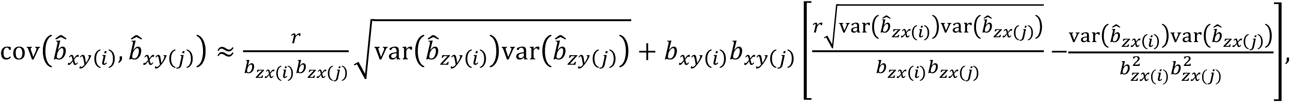
where subscripts *i* and *j* represent SNP *i* and *j*, respectively, *r* is LD correlation between the two SNPs (not available in the summary data but can be estimated from a reference sample with individual-level genotypes). The *i*-th diagonal element of **V** is
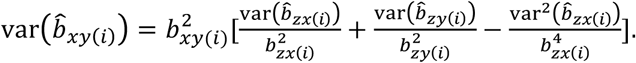
Therefore, we can estimate *b_xy_* from all the instruments using the generalized least squares approach as *b̂_xy_* = (**1**′**V**^−1^**1**)^−1^**1**′**V**^−1^**b̂**_*xy*_ with var (*b̂_xy_*) = (**1**′**V**^−1^**1**)^−1^. The statistical significance of *b̂_xy_* can be tested by *T_GSMR_* =
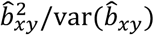
which follows a χ^2^ distribution with 1 degree of freedom.

### HEIDI-outlier: an approach to remove pleiotropic outliers

We have shown above that under a causal model the expected value of *b̂_xy_* estimated at any of the SNP instruments is identical in the absence of pleiotropy. If there are SNPs that have pleiotropic effects on *x* and *y*, *b̂_xy_* estimated at these SNPs will deviate from the expected value under a causal model, and hence will present as outliers. We previously proposed an approach (<u>he</u>terogeneity <u>i</u>n <u>d</u>ependent <u>i</u>nstrument, HEIDI) to test for heterogeneity in *b_xy_* estimated at multiple correlated instruments ^17^. Here, we extend this approach to detect heterogeneity in *b_xy_* estimated at *m* independent instruments (the method accounts for LD if there are remaining LD not removed by clumping) and then to remove outliers. The basic idea is to test where there is a significant difference between *b_xy_* estimated at an instrument *i* (i.e. *b*_*xy*(*i*)_) and *b_xy_* estimated at the SNP that shows the strongest association with exposure in the third quintile of the *b̂_xy_* distribution (i.e. *b*_*xy*(top)_). If we define *d_i_* = *b*_*xy*(*i*)_ - *b*_*xy*(top)_, we will have var (*d̂_i_* = var(*b̂*_*xy*(*i*)_ – *b̂*_*xy*(*top*)_) = var (*b̂*_*xy*(*i*)_) + var(*b̂*_*xy*(*top*)_ – 2cov(*b̂*_*xy*(*i*)_, *b̂*_*xy*(*top*)_), where cov(*b̂*_*xy*(*i*)_, *b̂*_*xy*(*top*)_) = 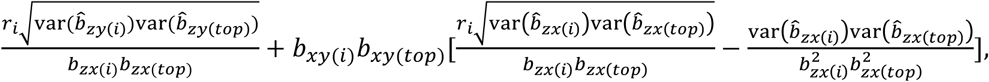
and *r* is the LD correlation between the two SNPs (estimated from a reference sample with individual-level genotypes). We can test the deviation of each SNP from the causal model using the χ^2^-statistic
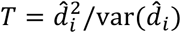
, and remove the SNPs with p-values < 0.01. We call this approach HEIDI-outlier. The number of tests involved in this analysis could be large if *m* is large. To retain as much power as possible to detect heterogeneity, we do not apply the stringent Bonferroni correction but use a modest threshold 0.01. Therefore, the ‘false positive rate’ is 0.01, meaning that even if there is no pleiotropic outlier, we will remove ~1% of the instruments, a very small proportion that is very unlikely to lead a substantial decrease of power in the subsequent GSMR analysis. We demonstrated by simulation that the difference in power with and without HEIDI-outlier filtering was subtle under a causal model without pleiotropy (**Supplementary Fig. 4a**). The test-statistics from real data analyses with HEIDI-outlier filtering were slightly smaller than that without HEIDI-outlier filtering (**Supplementary Fig. 4b**), suggesting that the analysis with HEIDI-outlier filtering is more conservative in terms of claiming causality.

### Multi-trait-based conditional GWAS analysis using summary data

To test whether the effect of a risk factor (*x*_0_) on a disease (*y*) depends on other risk factors (**x** = {*x*_1_, *x*_2_, ⋯, *x_t_*}), we usually perform a joint analysis based on the model below

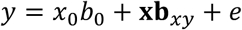

where *b*_0_ is the effect of *x*_0_ on *y*, **b**_*xy*_ = {*b_x_i_y_*} is a *t*-length vector with *b_x_i_y_* being the effect of a covariate *x_i_* on *y*, and *e* is the residual. Such an analysis is equivalent to a two-step analysis with the first step to adjust both *x*_0_ and *y* by **x** and the second step to estimate the effect of adjusted *x*_0_ on adjusted *y*. We therefore can estimate the effect size of *x*_0_ on *y* accounting for **x** by a GSMR analysis using SNP effects on *x*_0_ and *y* conditioning on **x**.

The conditional GWAS analysis usually requires individual-level genotype and phenotype data, which are not always available. Here, we propose a method to perform an approximate multitrait-based conditional GWAS analysis that only requires summary data. Since GWAS summary data for risk factors and disease are often from multiple independent studies, the analysis has to be performed conditioning on the genetic values of the covariate risk factors (denoted by **g**_*x*_ = {*g*_*x*_1__, *g*_*x*_2__,⋯, *g_x_t__*}). Following the method that uses GWAS summary data to perform a multi-SNP-based conditional and joint analysis (GCTA-COJO) ^60^, if we assume each covariate has been standardized with mean 0 and variance 1, the SNP effect on the disease accounting for **g**_*x*_ can be expressed as

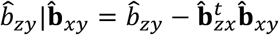

where *b̂_zy_* is the SNP effect on the disease on the logit scale (i.e. logOR), **b̂**_*xy*_ is a *t*-length vector with the *i*-th element *b̂_x_i_y_* being the effect of *g_x_* on the disease when all the covariates are fitted jointly ^28,47^, and **b̂**_*zx*_ is a *t*-length vector of SNP effects on **x**. We know from previous studies ^60^ that the joint effects of **g**_*x*_ on *y* (**b**_*xy*_) can be transformed from the marginal effects (**β**_*xy*_), i.e.

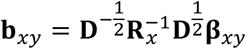

where **R**_*x*_ = {*r*_*g*(*x_i_*,*x_j_*)_} is a *t* × *t* matrix with *r*_*g*(*x_i_*,*x_j_*)_ being the genetic correlation between covariates *i* and *j*, **D** is a *t* × *t* diagonal matrix with the *i*-th diagonal element
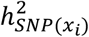
being the SNP-based heritability for the *i*-th covariate. We can estimate
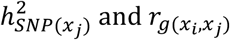
from GWAS summary data using the LDSC approaches ^28,47^, and estimate
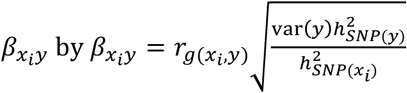
where
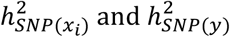
are the SNP-based heritability for *x_i_* and *y*, respectively, and *r*_*g*(*x_i_*,*y*)_ is the genetic correlation between *x_i_* and *y*. The variance of *y* on the logit scale can be estimated from the standard errors of the estimated logOR, i.e. var(*y*) ≈ var(*b̂_zy_*)2*p*(1 – *p*)*n* (**Supplementary Note**). We can estimate var(*y*) for each SNP and take the median value across all SNPs.

The sampling variance of *b̂_zy_*|**b̂**_*xy*_ is approximately

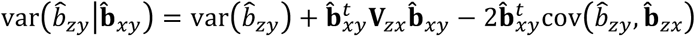

where **V**_*zx*_ = var(**b̂**_*zx*_), and cov(**b̂**_*zy*_, **b̂**_*zx*_) is a *t*-length vector with the *i*-th element cov(*b̂_zy_*, *b̂_zx_i__* being the covariance between *b̂_zy_* and *b̂_zx_i__*. We know from our previous study ^17^ that
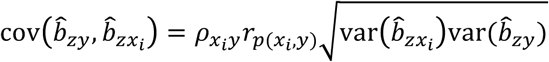
where *ρ_x_i_y_* is the proportion of sample overlap between *x_i_* and *y* and *r*_*p*(*x_i_*,*y*)_ is the phenotypic correlation between *x_i_* and *y*. In special cases, if *y* and **x** are observed in the same sample,
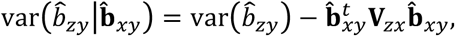
and if there is no sample overlap between *y* and **x**,
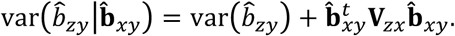
In practice, if there is a sample overlap between *y* and **x**, *ρ_x_i_y_ r*_*p*(*x_i_*,*y*)_ can be approximated by the intercept of a bivariate LDSC analysis between *x_i_* and *y* (ref ^28^). **V**_*zx*_ is the sampling variance-covariance of **b̂**_*zx*_ with the *ij*-th element
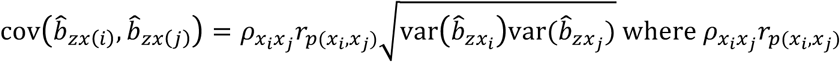
can also be approximated by the intercept of a bivariate LDSC analysis between *x_i_* and *x_j_*. The multi-trait-based conditional GWAS test can be performed using the test-statistic *T*_cond_ = (*b̂_zy_*|**b̂**_*xy*_)^2^/ var(*b̂_zy_* |**b̂**_*xy*_). We call this approach mtCOJO (<u>m</u>ulti-<u>t</u>rait-based <u>co</u>nditional and <u>jo</u>int analysis), and have demonstrated the accuracy of the approximation by simulation (Supplementary Fig. 5).

### GWAS data for risk factors and diseases

We used 9 risk factors as exposures for the GSMR analysis. These include 7 health risk factors i.e. body mass index (BMI), waist-to-hip ratio adjusted by BMI (WHRadjBMI), HDL cholesterol (HDL-c), LDL cholesterol (LDL-c), triglyceride (TG), systolic blood pressure (SBP) and diastolic blood pressure (DBP), and two additional phenotypes (height and educational attainment) that had a large number of instruments. We conducted GWAS analyses for SBP and DBP using data from the UK Biobank ^25^ (UKB) (see below for details of the UKB data). GWAS summary data for the other traits were from published studies (**Supplementary Table 2**). We re-calculated *b̂_zx_* from z-statistics using the method described in Zhu et al. ^17^ so that *b̂_zx_* could be interpreted in SD units. We then applied the clumping algorithm in PLINK ^26^ to select near-independent GWS SNPs for each trait (*r*^2^ threshold = 0.05, window size = 1Mb and p-value threshold = 5 × 10^−8^) using the 1000G-imputed ARIC data ^32^ (*n* = 7,703 unrelated individuals) as the reference for LD estimation. Since the statistical power of the GSMR analysis increases as the number of instruments, we performed the clumping analysis repeatedly for the SNPs in common between each pair of risk factor and disease data sets to maximize the number of instruments.

GWAS data for 22 common diseases were from two community-based studies. i.e., Genetic Epidemiology Research on Adult Health and Aging ^27^ (GERA) and UKB ^25^. There were 60,586 individuals of European ancestry in the GERA data. We cleaned the GERA genotype data using the standard quality control (QC) filters (excluding SNPs with missing rate ≥ 0.02, Hardy-Weinberg equilibrium test p-value ≤ 1 × 10^−6^ or minor allele count < 1, and removing individuals with missing rate ≥ 0.02), and imputed the genotype data to the 1000G using IMPUTE2 ^61^. We used GCTA ^62^ to estimate the genetic relationship matrix (GRM) of the individuals using a subset of the imputed SNPs (SNPs with minor allele frequency, MAF ≥ 0.01 and imputation INFO score ≥ 0.3 and in common with those in the HapMap phase 3, HM3), and computed the first 20 principal components (PCs) from the GRM. We removed one of each pair of individuals with estimated genetic relatedness ≥ 0.05 and retained 53,991 unrelated individuals for analysis. Individual-level ICD-9 codes were not available in dbGaP but had been classified into 22 common diseases (**Supplementary Table 3**). The disease status was coded as 0 (unaffected) and 1 (affected). We added an additional trait ‘Disease Count’ (a count of the number of diseases affecting each individual) as a crude measure of general health status of each individual. We then performed a genome-wide association analysis for each of the 23 phenotypes with age, gender and the first 20 PCs fitted as covariates.

Genotype data from UKB had been cleaned and imputed to a combined reference panel of 1000G and UK10K (see UKB documentation for details about QC and imputation). We included in the analysis only the individuals of European ancestry. Similarly as above, we computed the GRM and the first 20 PCs based on the HM3 SNPs with MAF ≥ 0.01 and imputation INFO score ≥ 0.3, and retained 108,039 unrelated individuals (GRM threshold of 0.05) for analysis. Individual-level ICD-10 codes were available in the UKB data. To match the diseases in GERA, we classified the phenotypes into 22 common diseases by projecting the ICD-10 codes to the classifications of ICD-9 codes in GERA taking into account self-reported disease status (Supplementary Table 3). We also added the trait ‘Disease Count’. We then conducted genome-wide association analyses for the 23 phenotypes using the same approach as above.

## URLs

GSMR R package: http://cnsgenomics.com/software/smr/gsmr.html

SMR: http://cnsgenomics.com/software/smr

PLINK: http://pngu.mgh.harvard.edu/~purcell/plink/

PLINK2: https://www.cog-genomics.org/plink2

GCTA: http://cnsgenomics.com/software/gcta

LDSC: https://github.com/bulik/ldsc

## Acknowledgments

This research was supported by the Australian Research Council (DP160101343), the Australian National Health and Medical Research Council (1107258, 1078901, 1078037, 1056929, 1048853 and 1113400), the Sylvia & Charles Viertel Charitable Foundation (Senior Medical Research Fellowship) and the Danish National Research Foundation (Niels Bohr Professorship). This study makes use of data from dbGaP (accession numbers: phs000090.v3.p1 and phs000674.v2.p2) and UK Biobank Resource (application number: 12514). A full list of acknowledgements to these data sets can be found in **Supplementary Note**.

